# Supersize me: torpor assists pre-hibernation fattening in a boreal bat

**DOI:** 10.1101/2024.05.23.595519

**Authors:** Mari A. Fjelldal, Niclas R. Fritzén, Kati M. Suominen, Thomas M. Lilley

## Abstract

Hibernators face an energetic dilemma in the autumn at northern latitudes; while temperatures and food availability decrease, hibernating species need to build fat deposits to survive the winter. During this critical fattening phase, insectivorous boreal bats use torpor to build and conserve their reserves. However, we still know little about temporal variability in torpor use employed by bats in autumn and how decreasing temperatures and food availability in combination with increasing individual body mass impact this. Here we present two general hypotheses for explaining temporal torpor patterns observed in a boreal bat (*Eptesicus nilssonii*), in which torpor use I) facilitates rapid mass gain or II) conserves stored body mass. Although temporally separated in our dataset, temperature, insect abundance and body mass throughout autumn in the study system indicate that *E. nilssonii* reaches a majority of its overwintering mass before the onset of increasing daily and nightly torpor use. In combination with generally low food availability by this point in time, these observations suggest that torpor expression might be intended to conserve gained reserves. Our study is intended as a first proof-of-concept for disentangling temporal drivers of torpor in bats during the pre-hibernation fattening phase.

## Introduction

All bat species at northern latitudes are insectivorous and therefore dependent on a food source that fluctuates heavily across the annual cycle (Speakman & Rowland 1999). During winter, when food is unavailable, bats hibernate and thus avoid some of the energetic challenges faced by other animals. However, winter hibernation requires a prior accumulation of large energy reserves if a hibernator is to survive until spring. Bats at high latitudes therefore face a critical pre-hibernation fattening phase, during which they must increase their body mass substantially (Kunz *et al*. 1998; Kronfeld-Schor *et al*. 2000). For insectivorous boreal bats, the building of fat deposits coincides with a time of decreasing temperatures and insect abundance (Speakman & Rowland 1999), adding to their challenge as the autumn progresses. Many hibernating bat species initiate the most rapid fat accumulation in late summer while temperatures and insect abundances are still relatively high (Kunz *et al*. 1998; Kronfeld-Schor *et al*. 2000; McGuire *et al*. 2009; Suba *et al*. 2010; Fraser & McGuire 2023). Although depositing fat stores early in the season might be more easily facilitated, it prolongs the duration that bats must preserve their large energy stores while elevating mechanical and metabolic power requirements to sustain foraging flights due to increased mass (Hughes & Rayner 1991). The timing and magnitude of fat deposition is therefore likely to be carefully managed within species and individuals.

To decrease energetic costs, bats can employ torpor across the annual cycle to lower metabolic requirements and thus conserve energy reserves (Stawski *et al*. 2014). The use of this strategy, i.e. the length and depth of the bouts, is however, highly impacted by environmental conditions (e.g. Wojciechowski *et al*. 2007), food availability (e.g. Coburn & Geiser 1998) and individual state (e.g. Fjelldal *et al*. 2023a), due to the dynamic balance of costs and benefits associated with the use of torpor (Boyles *et al*. 2020). During the pre-hibernation fattening period, bats rely on torpor to facilitate fat deposition (Speakman & Rowland 1999; McGuire *et al*. 2009; Suominen et al., *under review*). Still, until recently, torpor dynamics in bats during this critical phase have been unstudied. In the study by Suominen et al. (*under review*), torpor patterns in two bat species from the boreal zone were described throughout autumn for the first time, revealing strong temporal trends in torpor expressions, first increasing daily and then nightly torpor as the season advanced. However, the decline in temperature throughout the study period did not explain all of the variation observed in these torpor expressions, indicating that there were other temporally variable factors contributing to the strong shifts in strategic torpor management. By increasing torpor use during the daytime before initiating the use of torpor also at night, bats likely substantially reduce energetic requirements while still benefiting from foraging activities, thereby optimising their net energy gain. However, depending on the timing and intensity of the fat building phase, which can occur within a span of a few weeks (Kunz *et al*. 1998; Kronfeld-Schor *et al*. 2000), and the amount of insects available, the true benefit of this temporal and time-restricted specific torpor pattern is unknown.

Here, we present two hypotheses to explain both the increase in daytime torpor as well as the following increase in night-time torpor use, and relate these to food availability and increasing body mass from the end of the summer to the beginning of the winter. First, we hypothesize that peak mass gain is facilitated by maximising net energy gain through the initiation of increased use of daytime torpor at the onset of decreasing food availability, followed by an increase in use of night-time torpor after the peak mass gain (Fig. 1a). As an alternative, we hypothesize that body mass conservation after the peak mass gain is facilitated by the increased use of daytime torpor, and further, as foraging opportunities dwindle with further decreasing food availability, the increase in use of night-time torpor (Fig 1b). We will examine these hypotheses using empirical data on food availability, ambient temperature, and skin temperature and body mass of the boreal bat species *Eptesicus nilssonii*. Because our data does not provide a possibility to match body mass data to data on torpor use within individuals, and there are temporal discrepancies in data for food availability and mean temperatures, our goal is to provide an initial proof-of-concept with regards to the two presented hypotheses and fuel further investigation on this critical phase in the yearly cycle of insectivorous bats at high latitudes.

**Figure 1:**
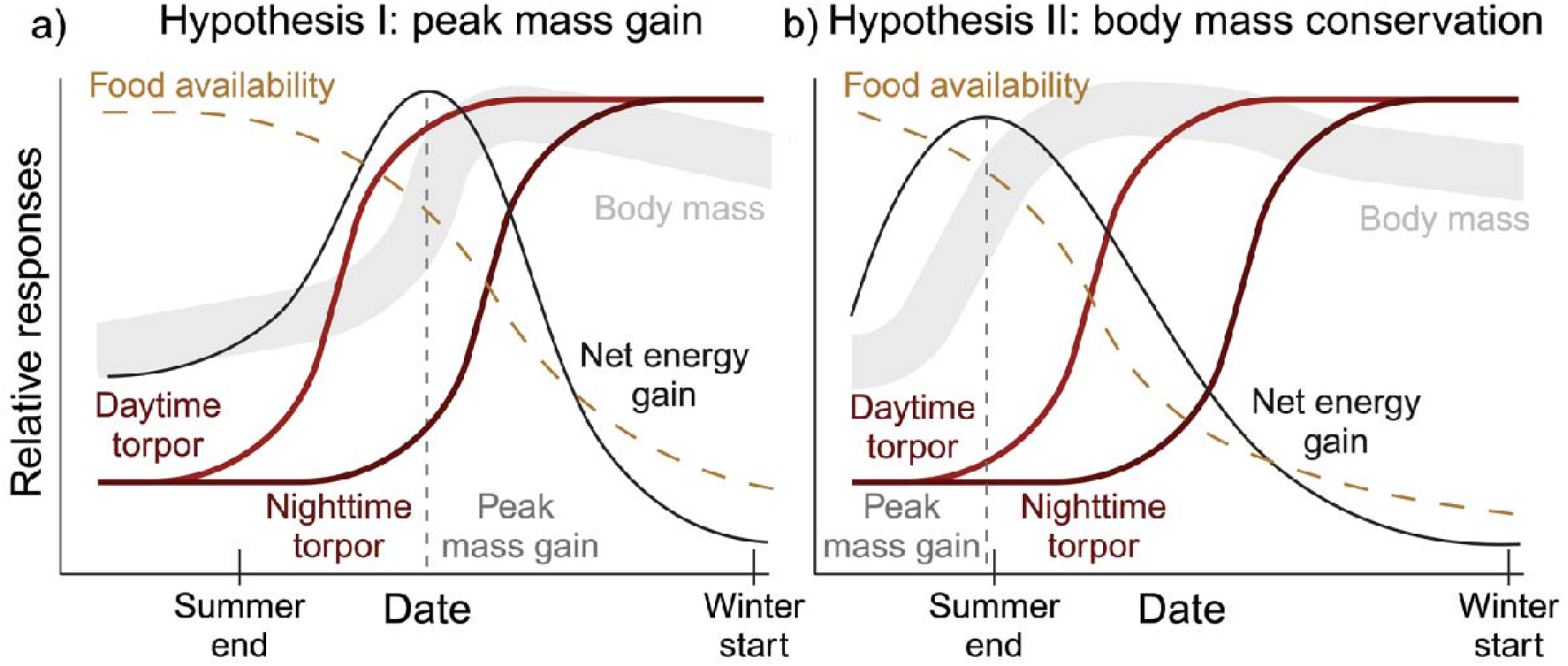
Graphic presentations of our two hypotheses for explaining the temporal torpor patterns observed in the study by Suominen et al. (*under review*). **a)** Hypothesis I describes the predicted relationships of food availability, net energy gain and body mass expected if the delayed increase in nightly torpor use (while increasing daily torpor) is intended for maximizing fat deposition. This would require high food availability as net energy gain reflects the energetic pay-off after accounting for metabolic and mechanistic costs of foraging flight. **b)** Hypothesis II assumes that the main body mass increase occurs in the late summer season, and that the increase in daily and nightly torpor use is expressed to conserve the already accumulated energy deposits as insect food diminishes.

## Results

In our high latitude study system (Valsörarna, Finland, 63°N), the empirical data collected during multiple years revealed distinct patterns for the environmental conditions throughout summer and autumn. Mean ambient temperature (based on data from 2014 to 2023) generally peaked at the end of July before declining steadily throughout August and September (Fig. 2a). However, insect productivity and -abundance (based on data from 2017 and 2018) begin to decline already in mid-July and reach relatively low levels by the beginning of September, although a single week in September 2018 showed a surge in insect counts (Fig. 2b).

**Figure 2:**
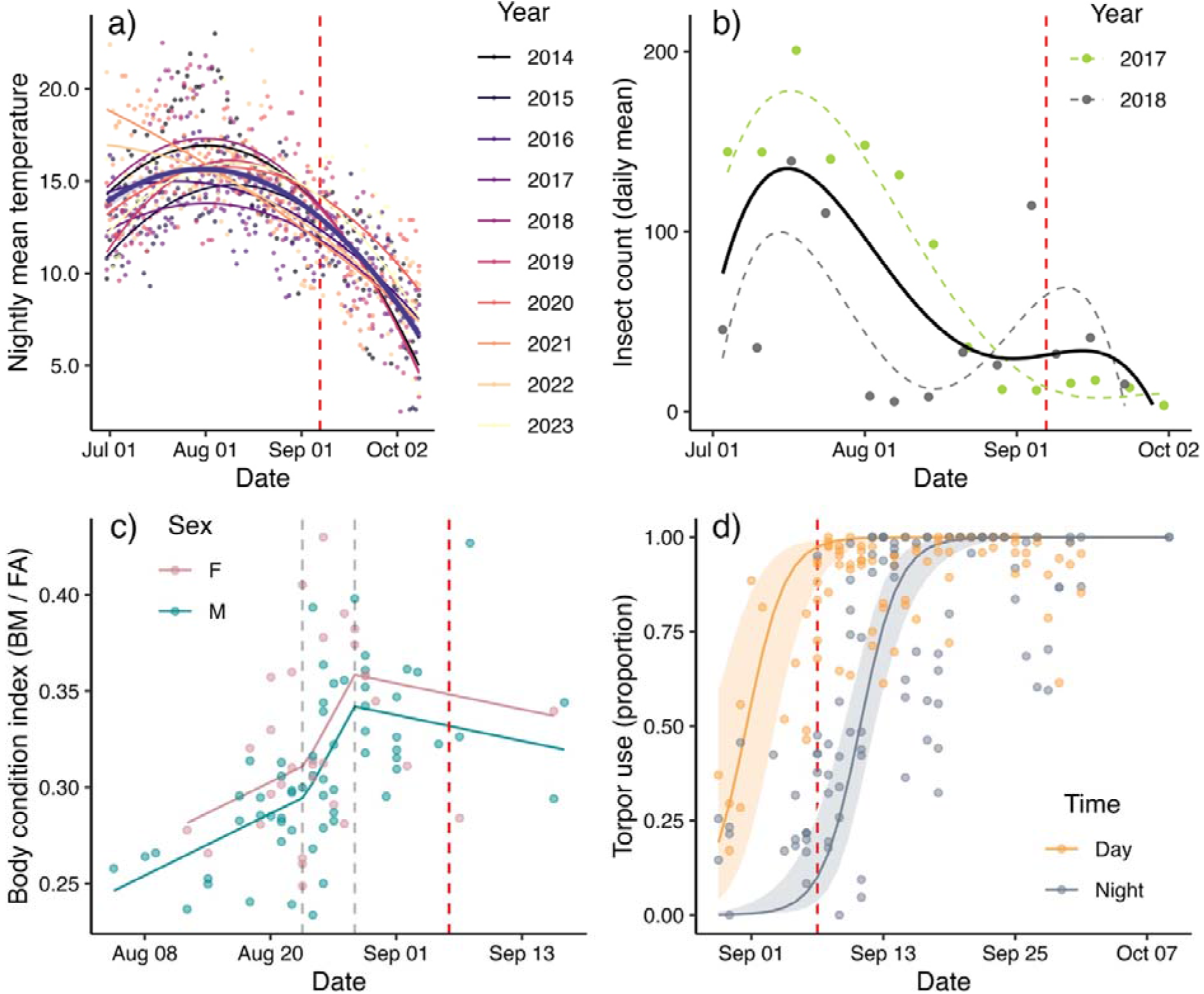
Temporal dynamics across the autumn for data collected during multiple years on the Valsörarna islands (Finland). Red dashed lines mark a common date (7^th^ September) for comparisons between panels. **a)** Nightly mean temperatures fitted as quadratic effects against date for each year (2014 to 2023), with the overall trend across years shown with the thicker dark line. **b)** Daily mean insect counts for each week during two consecutive years (grey and green) and the overall trend for both years (black) fitted as 4^th^ order polynomials against date. **c)** Body condition index (mass divided by forearm-length) of *E. nilssonii* against date, with males shown in blue and females in pink. Lines correspond to the predicted effects from the fitted breakpoint analysis with two identified break-points (Table S2); the first, indicating an increase in body condition, on the 24^th^ of August, and the second at the 29^th^ of August, indicating the predicted overall date of the peak body condition (dashed grey lines). **d)** The predicted logistic effect of torpor use (proportion) by *E. nilssonii* during day (yellow) and night (blue) in autumn, accounting for effects of wind and rain (Table S3).

Biometric data from adult *E. nilssonii* collected from 2014 to 2023 during captures in summer and autumn revealed strong temporal patterns in the overall body condition index. We identified two breakpoint dates in the data; the first detected at day 23.0 (24^th^ of August), where the increase in general body condition for males and females intensified after this point in time, until the second breakpoint on day 28.0 (29^th^ August), marking the overall date of the peak in predicted body condition of our dataset, after which the body condition index began to decline (Fig. 2c) (model results are shown in Table S1 and S2 in Supplementary Materials).

Torpor patterns for *E. nilssonii* in autumn were influenced by date, time of day (night versus day), rain and wind speed, as identified by Suominen et al. (*under review*) (model results are shown in Table S3 in Supplementary Materials). The date effect predicted a strong increase in daily torpor use by the end of August, until bats spent full days in torpor by early September (Fig. 2d). At this point in time, *E. nilssonii* also begun to increase nightly torpor use, reaching entire nights in torpor by mid-September (Fig. 2d).

## Discussion

Our results suggest the body mass conservation hypothesis (Fig. 1b) is better supported than the peak mass gain hypothesis (Fig. 1a) by the dataset available to us for the study. Although temporally disconnected, the dataset indicates the shift in torpor patterns (∼29^th^ August) occurs once *E. nilssonii* has passed the period of the peak mass gain (24^th^ – 29^th^ August), and as the decrease in food availability reaches a plateau. The increase in use of daily torpor and shift into using night-time torpor could represent a period of opportunistic feeding to maintain body mass dependent on individual state, environmental conditions and momentary food availability. Because bats in this study reach the majority of their overwintering mass already before the onset of shifts in torpor patterns, our dataset is not as supportive of the peak mass gain hypothesis, in which an increase in use of daily torpor and subsequent shift into using nighttime torpor would facilitate reaching maximum body mass.

Previous research on the use of torpor in the autumn has been inconclusive on the contributing drivers, with increasing Julian date being a stronger predictor than decreasing ambient temperatures (although correlated), suggesting that other temporal trends than temperature influence the torpor use observed in these bats during the pre-hibernation fattening (Suominen et al. *under review*). Here, we propose that the timing of increasing use of daytime torpor in *E. nilssonii*, followed by the increase in nighttime torpor, would be best explained by the combined effects of temporal changes to individual state and food availability, rather than just decreasing ambient temperature. It is notable that the breakpoint date (29^th^ August) detected for the timing of shifts in torpor use (Suominen et al. *under review*) corresponds to the breakpoint date of when maximum overwintering reserves have been reached. Although temporally disconnected, and therefore not available for investigation of direct associations, the data suggest the observed shift in torpor strategies from late summer to early autumn are triggered by the individual body condition in *E. nilssonii* reaching a certain level after a rapid peak mass gain period. However, this shift is most likely also influenced by food availability in the environment.

The diet of *E. nilssonii* has been found to consist of Nematocera, a large suborder of Diptera containing mosquitoes and midges (Vesterinen *et al*. 2018), in particular. Incidentally, Nematocera accounted for the majority of insects collected in this study (see Methods) and therefore, the temporal trend in insect decline described here is highly representative of the food availability for *E. nilssonii*, aligning with the predictions of the body mass conservation hypothesis (Fig. 1b). The stochasticity of food availability within and across years increases with an increase in latitude due to overall lower insect abundances (Pöyry *et al*. 2011). The unpredictable nature of food availability can contribute to a pattern in which maximum body mass is reached in advance of the ultimate decline in insect abundance, and switch to body mass conservation during a period when food availability is uncertain.

Although not evident in our system, the first hypothesis can also hold true for bats, depending on their feeding strategy. The early and rapid fattening, as observed here, appears to be mainly observed in ‘hawker’ bat species foraging on small aerial insects that decline rapidly in autumn, while ‘gleaner’ species, hunting terrestrial arthropod prey from the ground or vegetation surfaces, have food available for longer and can delay the onset or reduce the intensity of the lipogenesis (Reiter *et al*. 2010). In the study by Ignaczak *et al*. (2019), species such as *Myotis nattereri* and *M. bechsteinii*, with non-specialist diet composed of both aerial and terrestrial arthropods, appear to strongly delay the onset of pre-hibernation fattening or decrease their rate of mass gain. On the other hand, *Myotis daubentonii*, a hawker bat species hunting mainly aquatic insects, was observed to increase its body mass earlier and/or more rapidly than any of the gleaner bat species. These observations are similar to those recorded by Reiter *et al*. (2010) and support our overall hypothesis of food availability being one of the strongest drivers of species-specific pre-hibernation fattening strategies in insectivorous bats. We can therefore expect torpor patterns in autumn to also differ between species as a response to variability in expected foraging success and individual body condition throughout autumn.

Finally, we wish to highlight the need for empirical studies measuring the collective temporal trends of torpor use, food availability and body mass gain in bats during the critical pre-hibernation phase to better understand the interactions between environmental and individual conditions on thermoregulatory strategies. We see this initial foray in interpreting observed autumn torpor expressions in *E. nilssonii*, by investigating general temporal trends within the study system, as a basis for further investigation that can provide a more conclusive understanding on how hibernators can cope in a rapidly changing environment.

## Methods

The data presented in this study were collected on the Finnish Valsörarna islands (63°27’N, 21°46’E) located between Finland and Sweden in the outer archipelago of Kvarken in the Baltic Sea (Fig. 3). All handling and radio-tagging were carried out under licenses from the ELY Centres (EPOELY/1564/2023 & EPOELY/651/2023 and preceding licenses). All data was processed and analysed in R (version 4.3.1).

**Figure 3:**
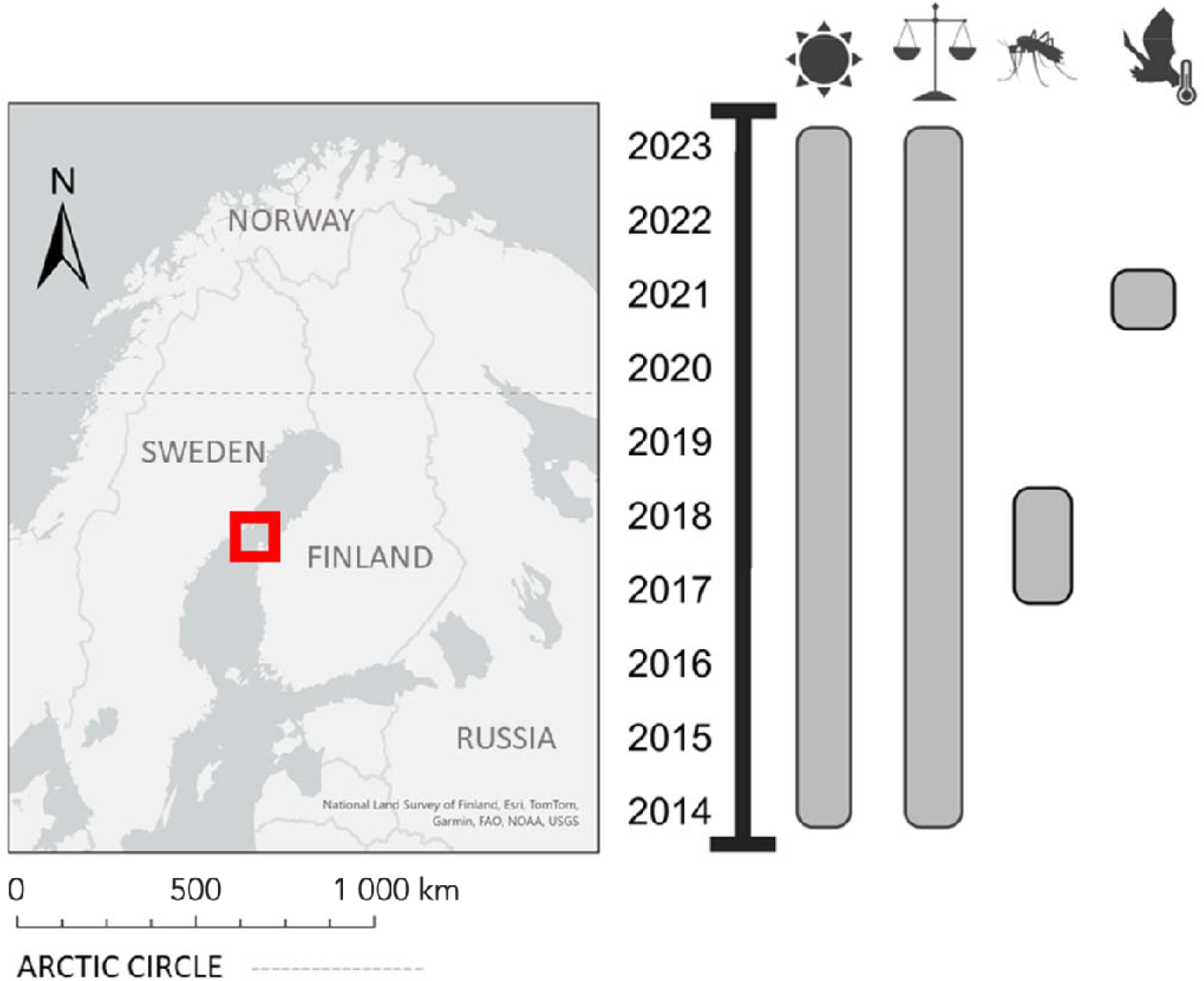
Map indicating location of study area (created with ArcGIS 3.3), and a timeline demonstrating the sampling years of air temperature, body weight measurements, insect collections, and skin temperature data of *E. nilssonii*, respectively.

### Body mass

The trapping of the bats was conducted within the KvarkenBats project maintained by the Valsörarna Biological Station. Bats were captured with harp-traps, mist-nets or during bat box checks between August and September each year from 2014 to 2023 (see Table S4 in Supplementary Materials for yearly capture dates). Information on sex, age class (adults versus juveniles), forearm-length and body mass were recorded for each individual. For the purpose of this study, we only considered the body mass of adult *E. nilssonii* (N_obs_ = 94). Initial linear models revealed a strong effect of forearm-length on body mass in our data, leading us to construct a body condition index (body mass (g) / forearm length (mm)), but no effect of time of capture (hours since sunset), and we thus excluded time of capture from further analyses. For this study we were interested in identifying dates indicating changes to the temporal trends in body condition throughout the study period. We therefore first performed a Davies’ test (library *segmented*) to test for the presence of breakpoints in a linear model with body condition index as the response and days since August 1^st^ and sex as explanatory variables. At least one breakpoint was detected (p = 0.019) with the best fit suggested around day 29.7. After confirming the presence of breakpoints, we tested for two breakpoints using the segmented function, with initial values set at 20 and 30, respectively, to identify and describe temporal shifts in the body condition index during the pre-hibernation fattening period.

### Torpor use

The torpor data presented here were first published in Suominen et al. (*under review*). Heat-sensitive radio transmitters (Telemetrie-Service Dessau, Telemetriesender V4 Temperatur) were attached to the back of captured *E. nilssonii* (N_ind_ = 12) using a skin glue (Sauer-Hautkleber). After releasing tagged individuals, pulse frequencies from each transmitter were recorded through an automatic radio-tracking system of ten RTEU-radio stations on the Valsörarna islands (see Gottwald *et al*. (2019) for details). We converted these pulse intervals to ten-minute skin temperature recordings and applied the methods described in Fjelldal *et al*. (2023b) to determine torpor bouts and -phases versus euthermic periods. In Suominen et al. (*under review*) a shift in temporal torpor patterns were discovered after conducting a breakpoint analysis on the data. Torpor before and after this breakpoint-date (29^th^ August) were analysed as separate periods, and for the purpose of this study we are only considering data after this breakpoint-date. The proportion spent torpid for each day (from sunrise to sunset) and night (from sunset to sunrise) was calculated and fitted against date in binomial generalized mixed models, with rainfall and wind speed as additional predictors and individual ID as random effect (see Suominen et al. (*under review*) for further details on analyses).

### Insect abundance

The collection of insects on the Valsörarna islands was initiated as part of a project for mapping the productivity of coastal lagoons and their importance to local ecosystems in the Kvarken area (Schneider & Fritzén 2020). Insects were collected throughout the summer and autumn in 2017 and 2018 using one malaise trap on land and two floating eclectors on the water surface (size ∼ 0.5 m^2^, at 10cm depth and at 30-50cm depth). All three traps were emptied on the same day once a week (after 5 to 9 days). The collected insects were preserved in 70% ethanol and counted based on order or sub-order; for the purpose of this study, we only considered insects belonging to the orders of Ephemeroptera, Lepidoptera (only moths), Nematocera, Neuroptera, Plecoptera, and Trichoptera. The Nematocera was by far the most abundant order, accounting for respectively 89% and 99% of the catches in 2017 and 2018. We merged the counts for all traps for each week and divided the count by the number of days insects had been collected. We only considered samples collected between 1^st^ of July until the last collection date each year (1^st^ October and 23^rd^ November, respectively). Total counts from these periods were 7729 insects in 2017 and 4234 insects in 2018.

### Temperature conditions

Temperature data throughout summer and autumn for each year from 2014 to 2023 were downloaded with ten-minute intervals from the meteorological station on the Valsörarna islands (Mustasaari Valassaaret), obtained through the Finnish Meteorological Institute.

## Acknowledgements

We thank the bat ringers Eeva-Maria Tidenberg, Ville Vasko, Jarmo Markkanen, Katarina Meramo, Anna Blomberg, Hanna Tuominen and the many other assistants and volunteers of the bat catching on the Valsörarna islands during the years 2014–2023. We also thank Michael Schneider for providing the insect data from the project Kvarken Flada.

## Funding statement

MAF was funded by Wihuri Foundation (grant number 00230067), NRF was funded by Waldemar von Frenckells stiftelse, KMS was funded by Kone foundation (grant number 201801142) and TML was funded by the Finnish Research Council (grant number 331515).

## Notes

### Competing Interest Statement

The authors have declared no competing interest.

