## Supplementary Materials for "Supersize me: torpor assists pre-hibernation fattening in a boreal bat"

**Model results for variation in body mass**

Results from the linear model used to identify breakpoints in the temporal trends in body condition are shown in Table S1. Results from the breakpoint analysis with two estimated breakpoints are shown in Table S2.

**Table S1**: Linear model results for explaining the variation observed in body condition index (mass / forearm-length) in *E. nilssonii*. Adjusted R^2^ = 0.23. *Note*: The effect of sex (males) is the difference from the intercept (females).

| Variable | Estimate (SE) | *p* value |
| --- | --- | --- |
| Intercept (female) | 0.26 (± 0.015) | < 0.001 |
| Sex (male) | -0.016 (± 0.0085) | 0.059 |
| Days since 1^st^ August | 0.0027 (± 0.0005) | < 0.001 |

**Table S2**: Results from the breakpoint analysis (function segmented) of the linear model presented above. Adjusted R^2^ = 0.31.

| Variable | Estimate (SE) | *p* value |
| --- | --- | --- |
| Intercept (female) | 0.25 (± 0.027) | < 0.001 |
| Sex (male) | -0.016 (± 0.0082) | < 0.05 |
| Days since 1^st^ August | 0.0027 (± 0.0013) | < 0.05 |
| BP1 (day 23.3) | 0.0072 (± 0.0074) | *NA* |
| BP2 (day 28.0) | -0.011 (± 0.0073) | *NA* |

**Model results for torpor use**

Results from the highest ranked model describing torpor use in autumn from the study by Suominen et al. (*under review*) are shown in Table S3. However, in this study we only consider data from the species *E. nilssonii*, and model values may therefore differ slightly from those presented in Suominen et al. (*under review*).

**Table S3**: Model results from the best logistic model identified in the study by Suominen et al. (*under review*), explaining variations in daily and nightly torpor use during autumn. *Note*: The nighttime effect is the difference in effect size when compared to the intercept (daytime torpor use).

|  | Random effects | Fixed effects | |
| --- | --- | --- | --- |
| Variable | Variance (SD) | Estimate (SE) | *p* value |
| *ID* | 4.5×10^-9^ (6.7×10^-5^) |  |  |
| Intercept (daytime) |  | -20.52 (± 4.21) | < 0.001 |
| Nighttime |  | -5.84 (± 1.22) | < 0.001 |
| Days since August 1^st^ |  | 0.56 (± 0.11) | < 0.001 |
| Mean wind (m/s) |  | 0.39 (± 0.14) | < 0.01 |
| Total rainfall (mm) |  | 0.50 (± 0.16) | < 0.01 |

**Yearly bat catches**

Sample sizes per year, species and sexes are shown in Table S4.

**Table S4**: Yearly captures of adult *E. nilssonii*.

| Year | First capture date | Last capture date | Males | Females | |
| --- | --- | --- | --- | --- | --- |
| 2014 | 26. August | 26. August | 1 | | 0 |
| 2015 | 30. August | 02. September | 1 | | 1 |
| 2016 | 26. August | 03. September | 6 | | 2 |
| 2017 | 21. August | 02. September | 8 | | 1 |
| 2018 | 21. August | 18. September | 5 | | 3 |
| 2019 | 19. August | 17. September | 13 | | 3 |
| 2020 | 18. August | 09. September | 6 | | 7 |
| 2021 | 09. August | 08. September | 9 | | 4 |
| 2022 | 13. August | 25. August | 6 | | 6 |
| 2023 | 06. August | 29. August | 6 | | 4 |
| **SUM** |  |  | **61** | | **31** |
